# Iterative computational bioprospecting of hydroxymethylfurfural oxidases combining molecular simulation and machine learning

**DOI:** 10.64898/2026.09.29.755274

**Authors:** Ruben Munoz-Tafalla, Mireia Martinez-Sugrañes, Juan Carro, Sergi Rodà, Victor Guallar, Francisco Javier Ruiz-Dueñas, Martin Floor

## Abstract

The enzymatic oxidation of 5-hydroxymethylfurfural (HMF) to 2,5-furandicarboxylic acid (FDCA) is a key step in the sustainable production of polyethylene furanoate (PEF), yet the final oxidation of 5-formyl-2-furancarboxylic acid (FFCA) to FDCA remains the principal bottleneck for industrial application. Although hydroxymethylfurfural oxidases (HMFOs) can catalyze this transformation, naturally occurring enzymes with high FFCA-oxidizing activity remain largely unexplored. Here, we present an iterative computational bioprospecting campaign to identify natural HMFOs with improved activity toward this rate-limiting reaction. Across four successive rounds, the computational workflow evolved from conservative homology-based screening to large-scale sequence mining assisted by machine learning, with each iteration redesigned according to the principal limitations identified through molecular simulations and experimental characterization. This adaptive strategy led to the discovery of several natural HMFOs with substantially improved FFCA oxidation activity, including candidates whose performance under the tested conditions approached that of the engineered benchmark 8BxHMFO. Beyond the identification of promising biocatalysts, the study demonstrates the value of continuous integration of computational prediction and experimental feedback for refining enzyme discovery campaigns, and provides practical insights into balancing catalytic prioritization, sequence diversity, developability, and computational scalability in future simulation-guided bioprospecting efforts.

**Graphical Abstract:** Iterative computational-experimental bioprospecting for natural hydroxymethylfurfural oxidases (HMFOs) with improved activity toward the oxidation of 5-formyl-2-furancarboxylic acid (FFCA) to 2,5-furandicarboxylic acid (FDCA). Computational predictions and experimental characterization were integrated across successive screening rounds, with experimental results guiding the redesign of subsequent computational strategies. This iterative approach progressively expanded the explored HMFO sequence space and led to the identification of natural enzymes with catalytic activities approaching those of highly active engineered variants.

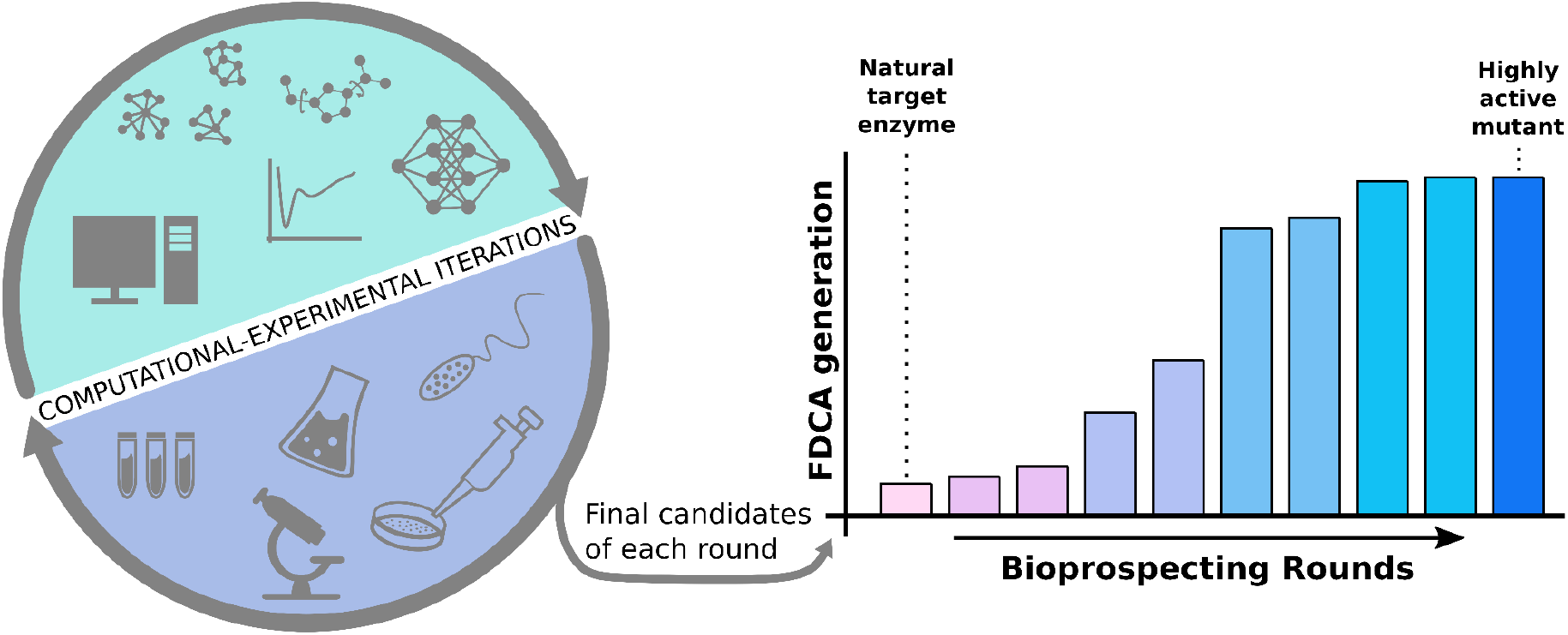

## Introduction

The transition from fossil-derived to bio-based chemicals is a central challenge in sustainable industrial chemistry. Among the most promising biomass-derived platform molecules, FDCA has emerged as a renewable replacement for terephthalic acid in polyester production. FDCA is the key monomer of PEF, a polymer that has attracted strong interest because of its improved barrier properties relative to polyethylene terephthalate (PET) and because of its potential role in lowering the fossil dependence of packaging and materials value chains (Bozell & Petersen, 2010; Gandini et al., 2016; van Putten et al., 2013).

FDCA is commonly obtained from HMF, which can be produced from biomass-derived sugars. The oxidation of HMF to FDCA proceeds through two main intermediates, 2,5-diformylfuran (DFF) and FFCA. Although chemical routes can achieve this transformation, they often require noble-metal catalysts and harsh operating conditions (Aranha & Gogate, 2023, Sheldon, 2018). Enzymatic oxidation offers an attractive alternative because it proceeds under mild conditions using molecular oxygen as the co-substrate and can provide high chemoselectivity (Vaidyanathan et al., 2023).

Hydroxymethylfurfural oxidases (HMFOs) are FAD-dependent members of the glucose-methanol-choline oxidoreductase superfamily that can catalyze the full HMF-to-FDCA oxidation cascade within a single active site. Their catalytic mechanism relies on a conserved His-Asn-His arrangement that helps position the substrate for hydride transfer to the flavin cofactor (Dijkman et al., 2015a). Despite this mechanistic suitability, the final FFCA-to-FDCA oxidation step remains the principal bottleneck for industrial application. Moreover, the number of HMFOs that have been experimentally characterized in detail remains limited, and most attention has focused on a small set of natural enzymes and engineered benchmark variants (Dijkman & Fraaije, 2014). Among these, the engineered variant 8BxHMFO, derived from the HMFO of Methylovorus sp. strain MP688 through the FRESCO protein engineering strategy, has emerged as a benchmark catalyst owing to its substantially improved thermostability and enhanced catalytic performance toward FFCA oxidation (Martin et al., 2018; Wijma et al., 2014).

This limited experimental landscape leaves open a central question for HMFO development: whether nature already contains diverse FFCA-competent catalysts that have not yet been sampled because discovery efforts have remained focused on a narrow neighborhood of known homologs or not. Answering this question is relevant not only for identifying improved catalysts, but also for understanding which properties should be prioritized during computational bioprospecting, including catalytic-pose stabilization, active-site diversity, phylogenetic diversity, and developability-related traits such as soluble expression and purification behavior.

Computational bioprospecting integrates sequence mining with structural modeling and molecular simulations to prioritize candidate enzymes before experimental characterization (Xiang et al., 2025). These approaches can substantially reduce the experimental search space while enabling exploration of enzyme diversity beyond the neighborhood of previously characterized homologs (Knox & Allen, 2023). More recently, machine learning has enabled the prioritization of candidates across much larger sequence spaces (Braun et al., 2023; Sampaio et al., 2023; Zhou & Huang, 2024). Although computational bioprospecting has become increasingly common for enzyme discovery, its application to HMFOs has remained limited.

Here, we present an iterative four-round computational bioprospecting campaign aimed at identifying natural HMFOs with improved FFCA oxidation activity. Starting from close homologs of experimentally characterized HMFOs, we progressively expanded the searchable sequence space while refining the computational prioritization strategy in response to experimental outcomes from each screening round. This iterative design-build-test-learn workflow ultimately enabled the identification of diverse natural HMFOs with strong FFCA oxidation activity while providing practical insights into the design of future computational enzyme discovery campaigns.

## Results and Discussion

### Round 1 - Conservative search into HMFO sequence space

The computational bioprospecting campaign began with a deliberately conservative exploration of the HMFO sequence landscape. The initial objective was to establish and validate a structure-based screening workflow within a region of sequence space where functional annotation was considered reliable. By restricting the search to close homologs of experimentally characterized HMFOs, uncertainties associated with structural modeling, active-site assignment, and mechanistic interpretation were minimized, providing a controlled starting point for the subsequent expansion of the campaign.

PSI-BLAST searches initiated from the three reference HMFOs retrieved 7,739 sequences from the UniProt/TrEMBL database. After removing sequences shorter than 400 amino acids and clustering at 80% sequence identity, a non-redundant dataset of 1,650 putative HMFOs was obtained. Construction of the sequence similarity network revealed a connected component comprising 36 closely related sequences at a 57% pairwise identity threshold (Figure 1A). This threshold was selected because it represented the highest pairwise identity cutoff that still produced a distinct connected cluster containing the three reference HMFOs while remaining separated from the rest of the network. These candidates were structurally prepared following the workflow described in the Methods section and evaluated using PELE simulations to estimate their Catalytic Binding Free Energy (CBFE) toward the FFCA oxidation step (figure 1B and 1C).

**Figure 1.**
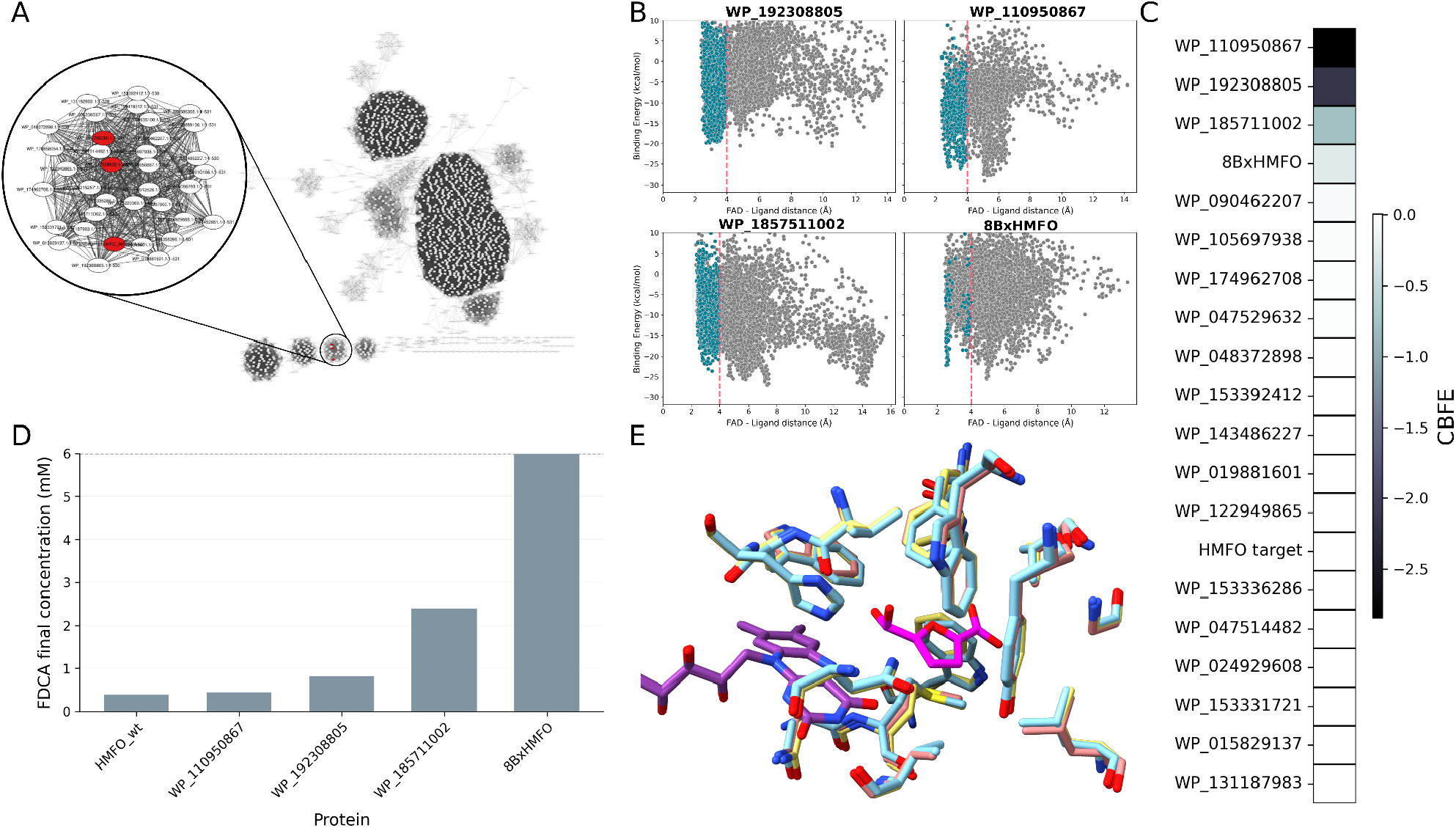
Computational and experimental results of round 1. A) SSN of the clustering of the 1650 non-redundant sequences at 57% identity threshold. The cluster of interest is amplified and search-seed sequences are colored In red. B) PELE results for the top 3 enzymes and 8BxHMFO. The poses of the whole simulation are colored in grey while poses accomplishing all catalytic distances, defined in the methods section, are colored in blue. C) Ranking of the candidates of the selected cluster based on CBFE. Color is based on the value of the calculated CBFE. being black the more negative values, and white the closer to 0 ones. D) Experimental results show the conversion from 6mM of FFCA to FDCA in 24 hours, with the following conditions 30°C, 200rpm. 1.5μM of enzyme, and 5μg/mL of catalase. E) Structural comparison of the active sites, where FAD cofactor is shown in purple and HFFCA ligand in pink. The three selected candidates residues closest to the ligand are shown in light blue, yellow, and light red, respectively.

The resulting CBFE distribution identified several candidates with predicted catalytic poses comparable to those of the reference enzymes. Three candidates, originating from Pseudomonas sp. PDM04 (WP_192308805), Pseudomonas bohemica (WP_110950867), and Pseudomonas cremoris (WP_185711002), were selected for experimental characterization on the basis of their computational ranking and sequence diversity.

All three enzymes were successfully expressed and showed measurable oxidative activity toward FFCA (Figure 1D). However, the overall conversion to FDCA remained limited. Under the tested conditions (6 mM FFCA, pH 8.0, 30 °C, 24 h), WP_185711002 was the most active candidate, producing approximately 30% FDCA conversion, whereas WP_110950867 and WP_192308805 generated only 0.4 and 0.9 mM FDCA, respectively. In contrast, when the full oxidative reaction was tested, all three enzymes efficiently oxidized HMF while accumulated FFCA as the predominant reaction product, confirming that the final FFCA-to-FDCA oxidation remained the principal catalytic bottleneck.

Although no improvement in FFCA oxidation was obtained during this first screening round, compared to the HMFO variant 8BxHMFO, the results provided an important proof of concept for the computational workflow. The PELE-based prioritization successfully identified catalytically competent HMFO homologs, even though the most active enzyme showed a less populated but narrower energy minimum during the simulation, suggesting a more defined interaction mode. The newly identified HMFOs also displayed highly similar active-site architectures to each other and to HMFO WT (Figure 1E), while the experimental characterization demonstrated that exploring only the immediate sequence neighborhood of known HMFOs was insufficient to uncover substantially improved natural catalysts. The limited functional diversity observed among these close homologs suggested that further progress would require expanding the search beyond conventional sequence-similarity criteria. This observation motivated the development of complementary candidate selection strategies focused on active-site diversification, described in the following section.

### Round 2 - Expanding the search through active-site diversification

The limited functional diversity observed among close homologs in Round 1 motivated the exploration of a broader region of HMFO sequence space. Rather than relying solely on global sequence similarity, candidate selection was redirected toward active-site diversity using two complementary strategies: clustering based on binding-site composition and design-guided natural mining (Figure 2A and 2B, respectively). Together, these approaches aimed to identify enzymes with distinct catalytic environments while preserving the canonical HMFO reaction mechanism.

**Figure 2.**
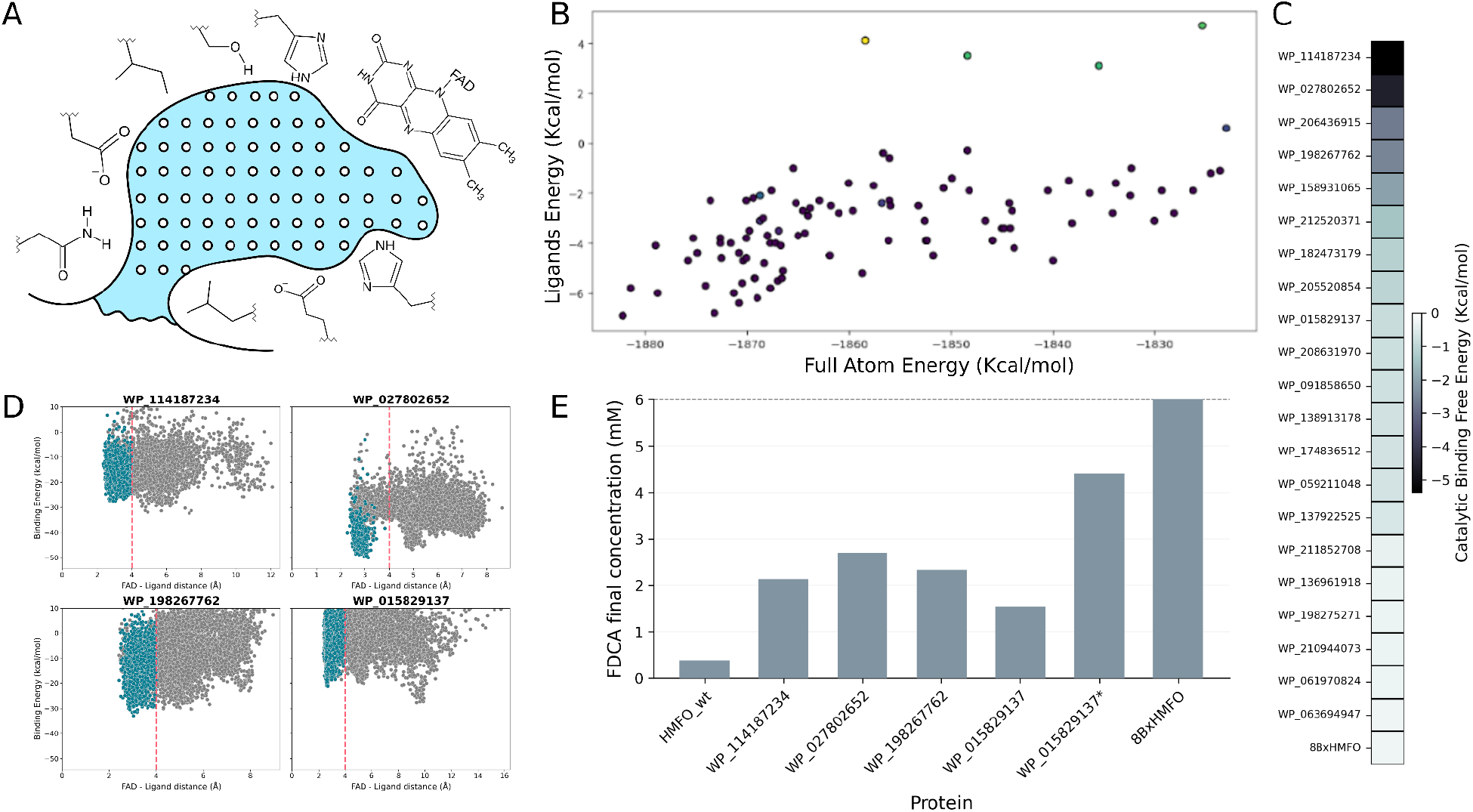
Computational and experimental results of round 2. A) Active site diversification approach using SiteMap: white spheres represent a volume unit, which is used to set the boundaries of the active site, and therefore calculate distances to the residues which delimit the cavity. B) Search for enhancing natural mutations approach using AsiteDesign. The plot shows the values of full atom and ligand’s energy for each mutant AsiteDesign generated. C) ranking of candidates of round 2 based on CBFE score. Color is based on the value of the calculated CBFE, being black the more negative values, and white the closer to 0 ones. D) PELE energy profiles of the experimentally tested enzymes. The poses of the whole simulation are colored in grey, while poses accomplishing all catalytic distances, defined in the methods section, are colored in blue. E) Experimental results of enzymes from round 2, compared to HMFO wt and 8BxHMFO variant, showing the conversion from FFCA to FDCA at a FFCA concentration of 6mM in 24 hours, with the following conditions: 30°C, 200rpm. 1.5μM of enzyme, and 5μg/mL of catalase. *Activity with FAD supplementation.

The first strategy focused on diversifying the composition of the substrate-binding pocket. Active-site residues were first defined on the reference HMFO from Methylovorus sp. strain MP688 using SiteMap. Two alternative pocket definitions were evaluated to assess the influence of residue selection on the recovered diversity. Clustering based on these active-site signatures revealed substantially greater variability than the homology-based selection used in Round 1 (Supplementary Figure S1). Representative sequences from each cluster were subsequently evaluated using PELE simulations.

In parallel, a design-guided natural mining strategy was implemented using AsiteDesign. Active-site substitutions predicted to improve interactions with FFCA in the reference HMFO scaffold were used as sequence signatures to screen the complete candidate dataset. This complementary strategy identified additional enzymes from sequence neighborhoods that were not recovered through active-site clustering alone, further expanding the diversity of candidates entering the computational screening pipeline.

Following PELE evaluation, candidates were prioritized for experimental consideration, from which the top ones were selected for heterologous expression (Figure 2C). Despite the improved catalytic predictions, experimental tractability emerged as a major bottleneck. Only one candidate, the HMFO from Methylovorus glucosotrophus (WP_015829137), the most similar to our starting target HMFO wt, was obtained in sufficient soluble yield for purification and detailed biochemical characterization. Three additional candidates, from *Microvirga aerophila* (WP_114187234), *Paraburkholderia dilworthii* (WP_027802652), and *Streptomyces sabulosicollis* (WP_198267762), were evaluated as soluble crude lysates after repeated attempts to improve soluble expression, including SUMO fusion and chaperone co-expression, failed to recover sufficient quantities of purified enzyme. Figure 2D shows the energy profiles for these 4 enzymes with HFFCA substrate. Even though candidate WP_027802652 shows a lower minima in binding energy, the top candidate in CBFE metric is WP_114187234, which has considerably more simulation steps in the correct conformation (blue circles).

Despite these experimental limitations, all four characterized enzymes displayed measurable FFCA oxidation activity (Figure 2E). This represented a clear improvement over the candidates identified in Round 1, which accumulated FFCA. WP_015829137 was the most active enzyme, producing 4.4 mM FDCA from 6 mM FFCA after 24 h at pH 7.0 and 30 °C. The enzyme, however, required supplementation with exogenous FAD to restore full activity following purification, indicating partial loss of the flavin cofactor during protein preparation. Among the enzymes evaluated as crude lysates, WP_027802652 and WP_198267762 each produced approximately 2.5 mM FDCA, whereas WP_114187234 reached 2.2 mM under the same conditions. As additional insights into their reaction mechanism, all four enzymes exhibited substrate inhibition at 15 mM FFCA, and only WP_027802652 showed detectable oxidation of the complete reaction cascade, producing approximately 1 mM FDCA after 24 h.

Collectively, these results demonstrated that expanding the search according to active-site composition successfully uncovered natural HMFOs with substantially improved FFCA oxidation activity compared with the close homologs identified in Round 1. At the same time, the relatively low success rate in soluble expression highlighted a new limitation of the campaign. By this stage, catalytic prioritization had become sufficiently effective that experimental developability, rather than the identification of catalytically promising sequences, emerged as the principal bottleneck. This observation motivated the incorporation of developability-oriented filters into the subsequent computational screening strategy.

### Round 3 - Large-scale computational bioprospecting

The improvements achieved in Round 2 demonstrated that expanding the search beyond close homologs could uncover natural HMFOs with enhanced FFCA oxidation activity. However, the limited number of candidates evaluated and the high rate of unsuccessful protein expression indicated that substantially broader exploration was required. Round 3 therefore aimed to scale the computational screening while maintaining sufficient diversity and experimental tractability to enable downstream characterization.

To maximize sequence diversity, filtering based on global sequence similarity was deliberately minimized. Starting from approximately 7,000 HMFO-like sequences containing the conserved catalytic His-Asn-His triad, candidates were subjected to the structural preparation and PELE workflow described in the Methods. In total, 1,570 enzymes were evaluated by PELE simulations, constituting the largest direct molecular simulation campaign performed in this study and the most computationally demanding stage of the bioprospecting workflow. The resulting CBFE values spanned a broad range, reflecting the increased structural and sequence diversity explored relative to the previous rounds. Figure 3A shows a sequence similarity network colored by CBFE from PELE.

**Figure 3.**
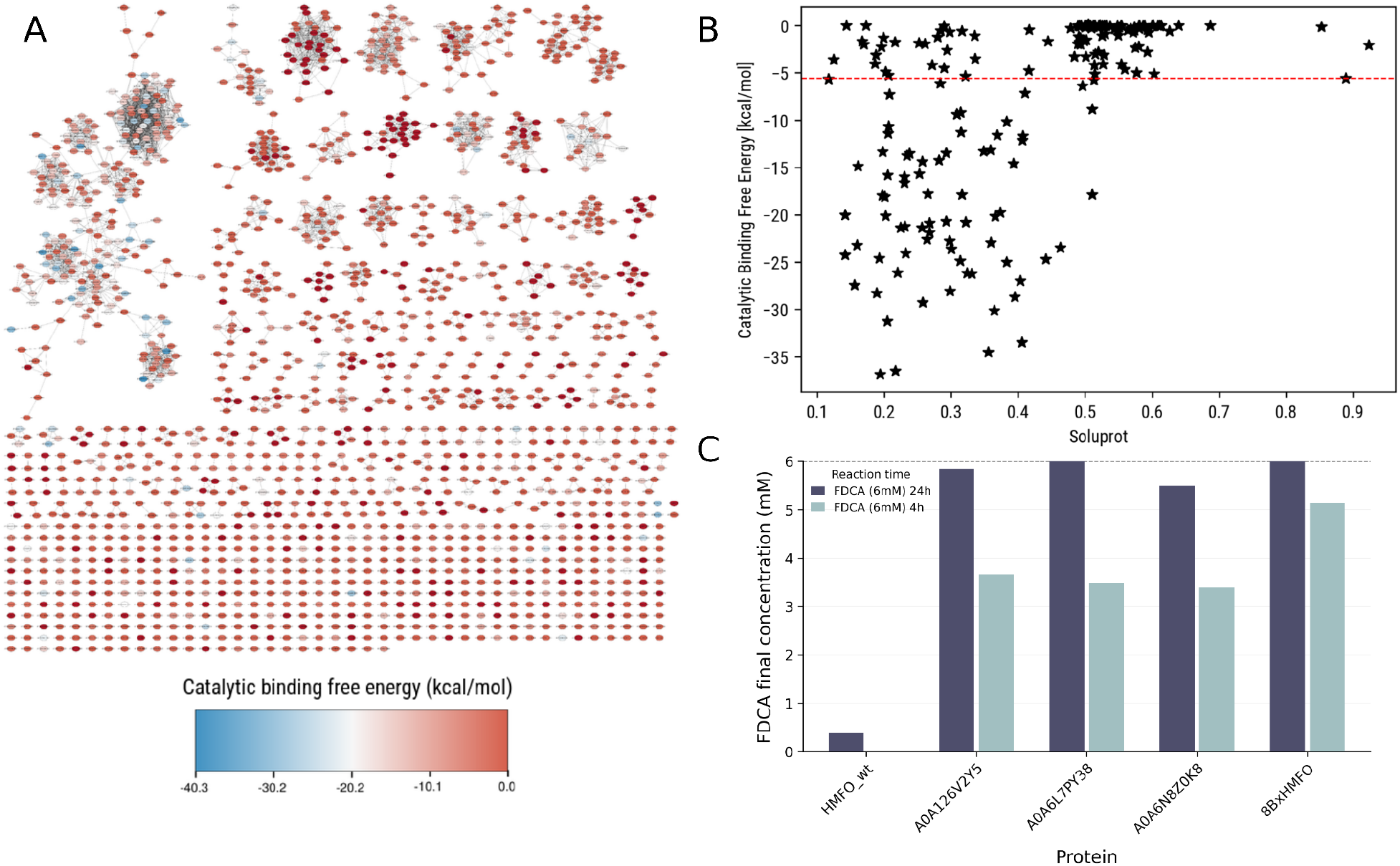
Computational and experimental results of round 3. A) SSN of the clustering of the newly found 1570 non-redundant sequences. CBFE value is used as the color, where more negative values of this metric are represented in blue, while values closer to 0 are shown in red. B) A comparative between CBFE and Soluprot solubility value of the simulated enzymes. C) Experimental results of the best 3 enzymes successfully expressed in round 3. Experimental results show the conversion from 6mM FFCA to FDCA in 24 hours and in 4 hours, compared to HMFO wt and 8BxHMFO variant, with the following conditions: 30°C. 200rpm. 1 5μM of enzyme, and 5μg/mL of catalase.

To improve the likelihood of successful experimental characterization, catalytic prioritization was complemented by developability assessment using SoluProt (Hon et al., 2021). Candidate sequences predicted to exhibit poor soluble expression in *E. coli* were discarded before selecting enzymes for experimental validation. This additional filtering step sought to reduce the experimental attrition observed in Round 2 while preserving the diversity recovered through the large-scale computational screening (Figure 3B).

The 14 highest-ranking candidates were subsequently selected for experimental characterization. Among them, only five enzymes (A0A286G251, A0A126V2Y5, A0A3D1LY36, A0A6L7PY38, and A0A6N8Z0K8) were successfully expressed in sufficient yield. Among these, A0A286G251 and A0A3D1LY36 showed no detectable activity toward the tested furanic substrates and were excluded from further analysis. In contrast, the remaining three enzymes exhibited a substantial improvement in FFCA oxidation compared with all candidates identified in the previous rounds (Figure 3C).

A0A6L7PY38 was the most active enzyme, achieving full conversion of FFCA to FDCA after 24 h at pH 8.0 and 30 °C. A0A126V2Y5 and A0A6N8Z0K8 also displayed high activity under the same conditions. All three enzymes exhibited little or no activity toward HMF or DFF, indicating that these enzymes remained specialized for the rate-limiting FFCA oxidation step.

Unlike the previous rounds, which primarily demonstrated the feasibility of the workflow and the value of increasing active-site diversity, the large-scale screening identified multiple natural HMFOs capable of approaching the FFCA oxidation performance of the engineered benchmark 8BxHMFO under the evaluated reaction conditions (Figure 3C). The combination of broad computational exploration with developability-oriented filtering therefore proved substantially more effective than either conservative homology searches or active-site diversification alone.

Although Round 3 markedly improved the efficiency of candidate discovery, the computational cost associated with performing PELE simulations at this scale became a limiting factor. In addition, the incorporation of the SoluProt developability predictor only partially alleviated the expression bottleneck identified in Round 2. While the proportion of successfully expressed candidates increased, unsuccessful heterologous expression remained a significant source of experimental attrition. Extending the same workflow to tens of thousands of candidate sequences would require prohibitive computational resources. This limitation motivated the incorporation of machine learning as a complementary prioritization strategy, enabling a much larger portion of HMFO sequence space to be explored while retaining the PELE-derived CBFE metric as the final criterion for structural evaluation.

### Round 4 - Machine learning-assisted expansion of the searchable sequence space

To overcome the high computational cost of testing a large amount of candidates, while significantly increasing the sequence space of search, the CBFE values generated during the previous round were used to train machine learning models capable of rapidly prioritizing candidate sequences directly from sequence-derived features, allowing a much larger portion of the HMFO sequence space to be explored.

Application of the trained models to the complete sequence dataset enabled the screening of 53,379 HMFO-like sequences, obtained after filtering 94,795 PSI-BLAST hits for conservation of the catalytic His-Asn-His motif (Figure 4A). Four independently trained models were used for candidate prioritization, each contributing its 300 highest-ranking sequences (Figure 4B). Following removal of duplicate entries and filtering using NetSolP expressibility predictions (Thumuluri et al., 2022), a final candidate pool comprising 300 unique enzymes was obtained. These candidates were subsequently subjected to the structural preparation and PELE workflow, with the resulting CBFE values serving as the final criterion for experimental prioritization. In parallel, a set of randomly selected sequences was also evaluated by PELE to provide a baseline for assessing the enrichment achieved by the machine learning strategy.

**Figure 4.**
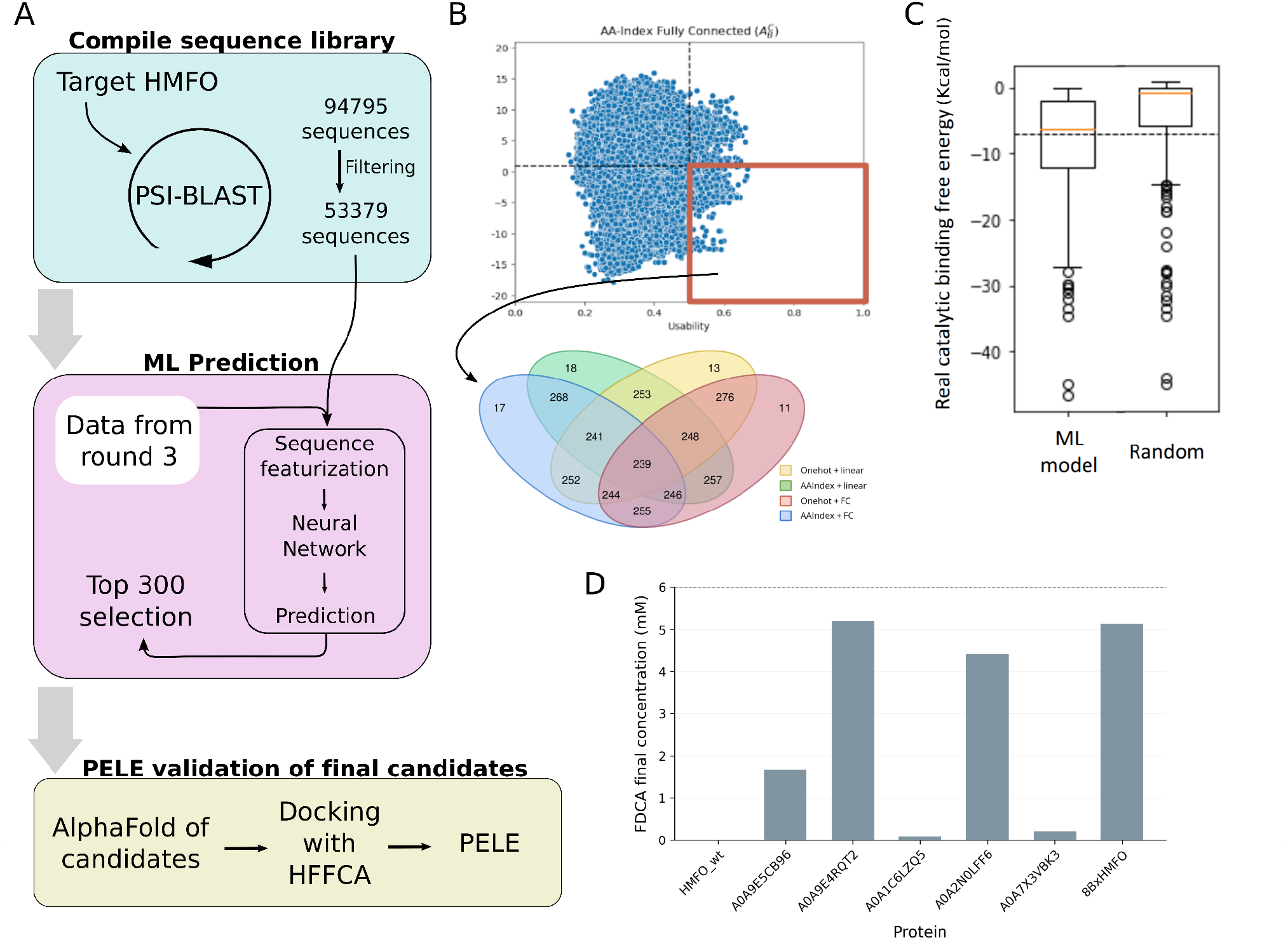
Computational and Experimental results of round 4. A) Workflow utilized to generate the sequence set the prediction of CBFE inferring results from round 3, and validation using PELE simulations. B) Selection of candidates based on predicted CBFE and predicted expressibility. The figure shows as an example the model A A-index Fully connected neural network. Sequence selection based on each of the four models can be seen below the plot. C) Comparison of PELE-derived CBFE values of ML-aided selection vs random selection of candidates. D) Experimental results of the successfully expressed enzymes in round 4. Experimental results show the conversion from 6mM FFCA to FDCA in 4 hours, compared to HMFO wt and 8BxHMFO variant, with the following conditions: 30°C, 200rpm, 1 5μM of enzyme. end 5μg/mL of catalase.

Comparison of the PELE-derived CBFE distributions demonstrated that the machine learning-guided selection substantially outperformed random sampling, enriching the candidate pool for enzymes predicted to stabilize catalytically competent binding poses (Figure 4C). These results indicate that the models successfully captured sequence patterns associated with favorable PELE-derived catalytic scores, thereby reducing the number of expensive molecular simulations required to identify promising candidates.

The top 34 candidates were selected for heterologous expression, of which eight enzymes were successfully characterized, including one candidate (A0A5N8YXA9) evaluated as a soluble crude lysate. Five of these enzymes (A0A9E5CB96, A0A9E4RQT2, A0A1C6LZQ5, A0A2N0LFF6, and A0A7X3VBK3) displayed measurable FFCA oxidation activity (figure 4D). Among all candidates identified throughout the study, A0A9E4RQT2, exhibited the highest FFCA oxidation activity. Under the evaluated reaction conditions, the enzyme achieved almost-complete conversion of 6 mM FFCA after 4 h at pH 8.0 and 30 °C. Its catalytic performance under these conditions was comparable to that of the engineered benchmark 8BxHMFO (Figure 4D).

Although the machine learning models successfully expanded the searchable sequence space and improved candidate prioritization, their predictive performance depended strongly on the representativeness of the available training data. The models were trained using the CBFE values obtained from the 1,570 candidates evaluated by PELE in Round 3, thereby leveraging a large, computationally expensive simulation dataset that had already been generated during the bioprospecting campaign. While this dataset provided an efficient foundation for model training, it was not necessarily representative of the full diversity of HMFO-like sequence space. Accordingly, prediction accuracy decreased for sequences that were increasingly divergent from those previously evaluated by PELE, indicating that the models primarily generalized within the sequence neighborhoods represented in the simulation-derived training dataset. Rather than replacing structure-based simulations, the results therefore support a complementary workflow in which machine learning efficiently identifies candidate-rich regions of sequence space, while PELE remains responsible for detailed structural evaluation and final prioritization.

Overall, Round 4 demonstrated that machine learning can substantially increase the scale of computational bioprospecting without replacing the underlying physics-based evaluation. By combining sequence-based prediction with targeted PELE simulations, the workflow successfully identified the most active natural FFCA oxidase discovered along all campaigns while substantially reducing the number of molecular simulations required to reach experimentally promising candidates.

### Broader implications and future directions

Although the computational bioprospecting campaign was primarily designed to identify natural HMFOs with improved activity toward the rate-limiting FFCA-to-FDCA oxidation step, the experimental characterization revealed substantial differences in substrate preferences among the enzymes identified across the different screening rounds. None of the individual enzymes efficiently catalyzed the complete HMF-to-FDCA oxidation sequence, but their distinct substrate preferences revealed complementary catalytic activities. In particular, A0A9E4RQT2 showed strong activity toward FFCA, whereas the Round 1 HMFO from Pseudomonas bohemica (WP_110950867) showed complementary activity toward the upstream oxidation of HMF and DFF. These complementary catalytic profiles provided a basis for combining naturally occurring HMFOs into a multienzyme cascade, demonstrating that the diversity recovered through computational bioprospecting can extend beyond the identification of individual high-performing catalysts.

A preliminary combination of WP_110950867 and A0A9E4RQT2 enabled complete conversion of 12.5 mM HMF to FDCA under the tested conditions, providing an initial demonstration that naturally occurring HMFOs identified through the campaign can be functionally integrated into a complete oxidation pathway. Further optimization of enzyme combinations, reaction conditions, and process configuration is beyond the scope of the present study, but would be valuable to explore in future work. More broadly, these observations suggest that future bioprospecting campaigns could consider complementary catalytic profiles alongside individual enzyme performance, particularly when the target transformation requires sequential reactions that may be inefficiently catalyzed by a single enzyme.

The broader outcome of the campaign is therefore not limited to the identification of individual HMFOs with improved FFCA oxidation activity. Across four successive rounds, experimental characterization was used to identify limitations in the preceding computational strategy and guide the design of the next iteration, while progressively expanding the explored sequence space. Notably, the experimental results obtained in each round showed an overall improvement over those of the preceding round, reflecting the progressive refinement of the computational strategy throughout the campaign (Figure 5). This iterative exchange between computational prediction and experimental observation enabled the workflow to evolve from targeted homolog screening to large-scale sequence mining assisted by machine learning. Importantly, all HMFOs identified in this study are naturally occurring enzymes without engineered mutations, including candidates whose FFCA oxidation performance approached that of the engineered 8BxHMFO benchmark. These enzymes therefore provide improved natural starting points for subsequent protein engineering, while their complementary catalytic properties open additional opportunities for the development of multienzyme systems.

**Figure 5.**
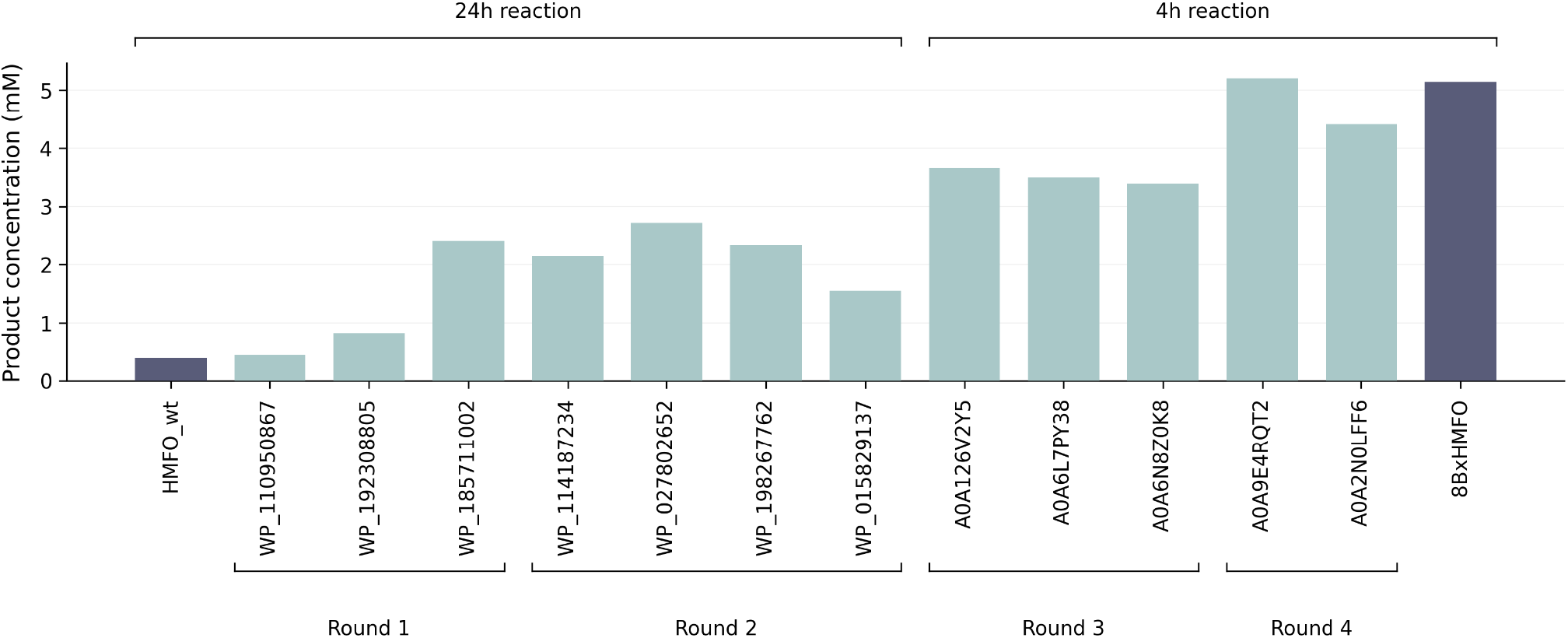
Activity of the top candidates identified in each screening round. HMFO WT Round 1, and Round 2 show FDCA formation after 24 h of reaction, whereas candidates from Round 3 and Round 4. as well as the 8BxHMFO variant, show FDCA concentration after 4 h of reaction Experimental conditions were 6mM substrate concentration, 30 °C, 200 rpm, 1.5 μM enzyme, and 5 μg/mL catalase.

## Conclusions

We present an iterative computational bioprospecting campaign to identify natural HMFOs capable of catalyzing the industrially relevant oxidation of FFCA to FDCA. Through four successive rounds of computational and experimental optimization, the workflow evolved from the exploration of close homologs to large-scale sequence mining assisted by machine learning, with each iteration addressing the principal limitation identified in the previous round. Crucially, experimental results were not treated solely as endpoint validation, but as feedback to guide the redesign of subsequent computational strategies, enabling the campaign to progressively improve candidate prioritization while adapting to newly identified bottlenecks. This approach led to the discovery of several natural HMFOs with substantially improved FFCA oxidation wild-type activity, including candidates whose performance under the tested conditions approached that of the engineered benchmark 8BxHMFO. More broadly, this study illustrates how iterative integration of computational prediction and experimental validation can provide practical insights that are transferable to future enzyme bioprospecting efforts beyond the HMFO family.

The results also clarify the complementary roles of the computational methodologies incorporated into this campaign. Structure-based molecular simulations proved effective for prioritizing candidates with a high likelihood of stabilizing catalytically relevant substrate-binding modes, while machine learning substantially expanded the searchable sequence space by leveraging the information generated through these simulations. At the same time, the campaign identified heterologous expression and model generalization as the principal remaining bottlenecks, highlighting that successful computational bioprospecting depends not on any single methodology, but on the successful iterative integration of complementary computational and experimental approaches.

## Acknowledgements

The authors gratefully acknowledge financial support from the Project FURENPOL (grant PLEC2021-007690), funded by MICIU/AEI/10.13039/501100011033 and by the European Union NextGenerationEU/PRTR, and Project ROBUSTOO, funded by the European Union (Grant Agreement No. 101135119).

## Author contributions

Conceptualization was led by V.G., and F.J.R-D. Computational methodology was developed by R.M-T., M.M-S., S.R., M.F., V.G., with R.M-T. responsible for the sequence-based and molecular simulation workflows and M.M-S. responsible for the machine learning framework. Experimental methodology was developed by J.C. and F.J.R-D. Investigation was carried out by R.M-T., M.M-S., S.R., M.F., and J.C. Supervision was provided by M.F., V.G., and F.J.R-D. Writing of the original draft was undertaken by R.M-T., M.M-S., M.F., V.G., and J.C., while review and editing were performed by R.M-T., M.M-S., J.C., S.R., M.F., V.G., and F.J.R-D.

## Data availability

Data from PELE simulations will be available in Zenodo in the future.

## Methods

### Sequence retrieval and clustering

Candidate HMFO sequences were identified by iterative PSI-BLAST searches against the UniProt/TrEMBL database using three experimentally characterized HMFOs as seed sequences: HMFO from Methylovorus sp. strain MP688 (NCBI WP_013440946.1), HMFO from Pseudomonas sp. 11/12A (NCBI WP_047529632.1), and HMFO from Pseudomonas nitroreducens (NCBI WP_024766380.1). Ten PSI-BLAST iterations were performed for each seed sequence. Retrieved sequences shorter than 400 amino acids were discarded to exclude truncated sequences and reduce the inclusion of unrelated short GMC oxidoreductases, and redundancy was reduced at 80% pairwise sequence identity using CD-HIT (Fu et al., 2012), resulting in a non-redundant dataset of 1,650 candidate HMFO sequences.

To characterize the sequence landscape, a sequence similarity network (SSN) was constructed from all-against-all BLASTP pairwise sequence identities. The similarity matrix was converted into an undirected graph using Cytoscape (Shannon et al., 2003), where nodes represented protein sequences and edges connected sequence pairs with pairwise identities above a specified threshold. For the initial screening round, a threshold of 57% pairwise identity was applied, yielding a connected component comprising 36 sequences, including the three reference HMFOs. This subset served as the starting point for the first computational screening campaign.

### Structural modeling and protein preparation

Structural models of all selected HMFO candidates were generated using AlphaFold2 (Jumper et al., 2021). Prior to structural preparation, the presence of the canonical catalytic His-Asn-His triad was verified by multiple sequence alignment with the three reference HMFOs. Candidates lacking conservation of the three catalytic residues at the corresponding alignment positions were excluded from subsequent analyses.

To improve model quality for molecular simulations, disordered terminal regions were removed based on AlphaFold pLDDT confidence scores. The trimmed structures were aligned to the crystal structure of HMFO from Methylovorus sp. strain MP688 (PDB ID: 4UDP), which served as the structural reference throughout the study. The FAD cofactor and crystallographic active-site water molecules were transferred from the reference structure to each AlphaFold model by structural alignment.

Prepared models were subsequently refined using the FastRelax protocol implemented in Rosetta (Alford et al., 2017), with restraints applied to preserve the geometry of the active site while relieving local structural strain introduced during model preparation. Following refinement, the hydrated form of 5-formyl-2-furancarboxylic acid (HFFCA) was docked into the active site using Glide from Schrödinger (Friesner et al., 2004) to generate the initial enzyme-substrate complex for subsequent PELE simulations. The geometry of the protein-cofactor-substrate complex was inspected prior to production simulations to verify the integrity of the catalytic site.

### PELE simulations and Catalytic Binding Free Energy (CBFE)

Enzyme-substrate interactions were evaluated using Protein Energy Landscape Exploration (PELE), a Monte Carlo-based molecular simulation framework that enables adaptive sampling of ligand binding and protein conformational changes (Borrelli et al., 2005; Lecina et al., 2017). All simulations were performed on the MareNostrum 4 supercomputer at the Barcelona Supercomputing Center. Each PELE simulation consisted of 56 independent trajectories. Every trajectory included 100 equilibration steps followed by 300 PELE steps each. The prepared enzyme-FAD-HFFCA complexes described above were used as the starting structures for all simulations.

To identify catalytically relevant binding modes, simulation snapshots were evaluated against four geometric criteria defining a near-attack conformation (NAC): (i) the distance between the hydride-donor hydrogen of HFFCA and the N5 atom of the FAD cofactor (≤ 3.5 Å); (ii) the distance between the transferable hydroxyl hydrogen of HFFCA and the NE2 atom of the catalytic histidine (≤ 3.5 Å); (iii) the distance between the corresponding hydroxyl oxygen of HFFCA and the side-chain oxygen of the catalytic asparagine (≤ 3.5 Å); and (iv) the distance between the ND1 atom of the proton-relay histidine and the HD1 hydrogen of the catalytic histidine (≤ 3.5 Å). Because HFFCA contains two hydroxyl groups that may participate in the oxidation reaction, criteria (ii) and (iii) were evaluated for both possible donor–acceptor assignments. Only geometrically consistent assignments, in which the transferable hydrogen was covalently linked to the interacting oxygen atom, were considered. Simulation snapshots satisfying all four geometric criteria simultaneously were classified as catalytically competent.

For each enzyme, a Catalytic Binding Free Energy (CBFE) value was calculated as the Boltzmann-weighted average binding energy of all catalytically competent snapshots, using this formula:

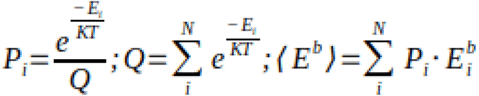

Where Q is the partition function for a simulation consisting of N poses (in this case, only catalytic poses are considered for the calculation). Eb represents the binding free energy, which is calculated by weighting the probability of each pose by its interaction energy obtained from PELE. Throughout this study, CBFE was used exclusively as a relative prioritization metric to rank candidates according to their ability to stabilize catalytically competent binding poses, rather than as a direct predictor of enzymatic activity or turnover.

### Binding-site analysis and diversification

To expand the structural diversity explored beyond close homologs of the reference HMFOs, candidate enzymes were compared according to the composition of their substrate-binding pockets rather than overall sequence similarity. The catalytic pocket of the reference HMFO from Methylovorus sp. strain MP688 was identified using SiteMap from Schrödinger (Halgren, 2009) with a pocket-point distance threshold of 2.5 Å, a site box size of 10 Å, and fine grid resolution.

Two alternative definitions of the binding site were evaluated. In the first, all residues containing at least one atom within 2.5 Å of the SiteMap pocket volume were considered part of the binding site. In the second, only residues whose side chains were oriented toward the pocket, defined as side-chain atoms located within 2.5 Å of the SiteMap volume points, were retained. These two definitions were intended to evaluate whether excluding solvent-facing or structurally peripheral residues improved the identification of functionally relevant active-site variation.

For each binding-site definition, the corresponding active-site sequences were extracted from a multiple sequence alignment of the 1,650 non-redundant HMFO candidates. Pairwise similarities between binding-site sequences were calculated using the BLOSUM62 substitution matrix, and candidates were clustered into 50 groups according to their active-site similarity. Representative sequences were selected from each cluster for subsequent structural preparation and PELE evaluation.

### Design-guided sequence mining

To complement the binding-site diversification strategy, a design-guided approach was employed to identify naturally occurring HMFOs carrying active-site substitutions predicted to improve substrate binding. Active-site redesign was performed on the reference HMFO scaffold using AsiteDesign (Roda et al., 2023), allowing substitutions at selected binding-site positions while maintaining the catalytic residues fixed. The resulting designs were analyzed to identify amino acid substitutions that were consistently enriched among the highest-scoring variants.

The non-redundant HMFO sequence dataset was subsequently screened for naturally occurring enzymes containing these predicted beneficial substitutions. Candidate sequences carrying at least two of the identified substitutions were selected for further evaluation. Structural models of the selected enzymes were retrieved from the AlphaFold Protein Structure Database, prepared as described and evaluated using the PELE simulation protocol described in the corresponding methods sections.

### Developability assessment

As the scale of the computational screening increased, developability predictors were incorporated into the candidate prioritization workflow to reduce experimental attrition arising from poor heterologous expression. Solubility predictions were used as an additional filtering criterion prior to experimental characterization, complementing the structural and catalytic prioritization provided by PELE.

Two prediction tools were employed. SoluProt is used to estimate the probability of soluble recombinant expression in Escherichia coli based on sequence-derived features (Hon et al., 2021). NetSolP is used to predict both protein solubility and overall usability for recombinant expression (Magnan et al., 2024). Candidate sequences were prioritized according to their predicted developability scores, and, where applicable, only sequences exceeding the selected prediction threshold were retained for downstream evaluation.

Developability predictions were used exclusively as a pre-filter to improve the likelihood of obtaining soluble recombinant proteins and were not considered indicators of catalytic performance. Final candidate selection was therefore based on the combined assessment of predicted developability and PELE-derived CBFE values.

### Machine learning-assisted prioritization

#### Training dataset and prediction targets

To extend computational bioprospecting beyond the number of candidates that could be directly evaluated by PELE simulations, a supervised machine-learning framework was developed to predict simulation-derived catalytic metrics directly from protein sequence. The objective of the models was not to predict experimentally measured catalytic activity, but rather to reproduce the sequence-to-score relationship obtained from the PELE simulations and thereby provide a computationally inexpensive pre-screening step before explicit structural modelling and simulation.

The ML dataset was derived from the large-scale HMFO bioprospecting campaign described above. Starting from the HMFO sequence library, candidates were clustered using an 80% maximum pairwise sequence identity threshold, structurally modelled and evaluated using the PELE workflow. The final simulation-derived dataset contained approximately 1,570 HMFO homologs for which catalytic PELE trajectories and the corresponding energetic descriptors were available.

Two simulation-derived metrics were initially considered. First, the mean of the ten lowest binding energies among poses fulfilling the catalytic geometric criteria. Second, the Catalytic Binding Free Energy (CBFE), corresponding to the Boltzmann-weighted binding score calculated over the ensemble of catalytically competent poses. Both metrics were benchmarked during model development. Because the CBFE incorporates information from the probability distribution of the sampled catalytic binding-energy landscape, it was subsequently retained as the principal metric for the large-scale Round 4 bioprospecting application.

#### Target normalization

The distribution of the PELE-derived target values was examined before model training because the two energetic metrics showed markedly different degrees of skewness. Two normalization schemes were therefore evaluated. In the first, the target values were standardized using Z-scores. In the second, Z-score normalization was followed by a cubic-root transformation to reduce skewness in strongly asymmetric target distributions.

The effect of the normalization strategy was evaluated independently for each PELE-derived metric during model benchmarking. In particular, the CBFE distribution was strongly skewed toward unfavorable values, with comparatively few sequences populating the favorable energetic region.. The combined Z-score and cubic-root transformation substantially reduced this asymmetry and was consequently used for the CBFE-based models selected for the large-scale bioprospecting stage.

#### Sequence feature representations

Various alternative protein-sequence representations were evaluated to determine how sequence encoding affected the ability of the models to reproduce the PELE-derived metrics. For alignment-based representations, all protein sequences were aligned using MAFFT (Katoh et al., 2002) and converted into numerical feature vectors using one-hot encoding, AAindex-based physicochemical encoding (Kawashima et al., 2008), or a concatenated representation combining both approaches.

In the one-hot representation, each aligned position was encoded using a 21-dimensional categorical vector corresponding to the 20 standard amino acids and an additional gap state, and the resulting vectors were concatenated across the alignment. For the physicochemical representation, amino-acid properties were obtained from the AAindex database. The initial 566 descriptors were reduced to 19 components by principal-component analysis, and the resulting residue-level vectors were concatenated across the alignment.

As an additional control during model development, random residue encodings of the same dimensionality as the reduced AAindex representation were generated to evaluate whether the performance associated with the AAindex models resulted from chemically meaningful amino-acid information rather than simply from an increased numerical dimensionality of the input.

Protein language-model representations were also investigated. Sequence embeddings were generated using pretrained ESM-2 protein language models (Rives et al., 2021; Lin et al., 2023). The principal model used for benchmarking was ESM2_T33_650M_UR50D, while the larger ESM2_T36_3B_UR50D model was additionally evaluated to assess whether increasing language-model capacity improved prediction of the PELE-derived quantities.

#### Machine-learning architectures

Two supervised regression architectures of different complexity were investigated: a linear regression model and a fully connected neural network. The linear model consisted of a single fully connected output layer connecting the complete input feature vector directly to one scalar prediction, whereas the neural network comprised an input layer with 1,280 units, a hidden fully connected layer with 1,024 units, and a single-unit output layer. Leaky rectified linear unit activation functions were used in the hidden layer. The models were implemented using Python 3.8 and PyTorch 2.0.1 (Paszke et al., 2019).

#### Model training and cross-validation

Model performance was assessed using 10-fold cross-validation. For each fold, 10% of the complete simulation-derived dataset was retained as an independent test partition, while the remaining 90% was divided into training and validation subsets using an 80:20 ratio. The training partition was used for parameter optimization and the validation partition for model development and hyperparameter selection; the test partition was kept separate and used to assess performance on unseen sequences.

Models were optimized using the Adam optimizer and mean-squared error (MSE) as the loss function (Kingma & Ba., 2015). Training was continued until convergence, with the model showing the best validation performance retained. To quantify variability arising from random partitioning of the sequence dataset, 100 independent replicas were performed for each model configuration. Model training was carried out using CPUs on the Nord III cluster at the Barcelona Supercomputing Center.

Predictive performance was evaluated using Pearson and Spearman correlation coefficients to quantify linear and rank-order agreement, respectively, between predicted and PELE-derived values. Because the practical objective was to enrich for favorable candidates rather than reproduce the complete target distribution, model performance was additionally assessed using a ranking-based evaluation. For each test partition, sequences were ranked according to their predicted score, and the best PELE-derived value recovered within increasing candidate-selection budgets was recorded. This metric quantified the ability of each model to recover highly ranked candidates from a restricted prediction set and was used together with the correlation metrics to select models for the Round 4 screening campaign.

#### Influence of training-set size

The amount of simulation-derived data required for useful predictive performance was evaluated by repeating the training procedure with progressively smaller subsets of the available training data. The original 90% train-validation partition was subsampled to maximum dataset sizes of 10, 50, 100, 200, 500, 750, 1,000 and 1,250 sequences, with each subset divided into training and validation data using the same 80:20 ratio as in the main training protocol.

The analysis was performed using four representative model configurations: linear and fully connected models trained with either one-hot or AAindex features. Both PELE-derived target metrics were evaluated to determine whether useful sequence-to-score models could be obtained with substantially fewer PELE-labelled sequences.

#### Large-scale ML-assisted HMFO screening

Following benchmarking, four model configurations were selected for the last bioprospecting iteration: linear regression and fully connected models using either one-hot or AAindex sequence encoding. These models showed useful performance in the CBFE ranking task while covering both sequence representations and model architectures.

A larger HMFO-like sequence collection was generated by performing PSI-BLAST searches against the UniProt/TrEMBL database (Bairoch et al., 2000) using the wild-type HMFO from Methylovorus sp. strain MP688 as the query sequence. No maximum number of retrieved hits was imposed, yielding 94,795 sequences. Candidates were filtered for conservation of the catalytic His-Asn-His triad, resulting in 53,379 sequences for ML prediction.

Because low soluble-expression yields had been an important source of attrition in previous rounds, protein developability was incorporated using NetSolP (Thumuluri et al., 2022). Only sequences with a predicted usability score >0.5 were considered for downstream prioritization. This metric incorporates predicted expression and solubility and was used exclusively as a developability filter rather than as a surrogate for catalytic performance.

The four selected models independently predicted CBFE values for the 53,379 retained sequences. Candidates were ranked after applying the NetSolP cutoff and requiring a predicted CBFE more favorable than that of the engineered HMFO benchmark. For each model, the 300 highest-ranking sequences satisfying these criteria were retained. The resulting lists were merged and duplicate sequences removed before structural evaluation.

ML predictions were used exclusively as a computational pre-screen. The resulting candidates were therefore returned to the physics-based workflow and evaluated using the same structural preparation, substrate docking, and PELE simulation procedures described in Sections 2.2 and 2.3. The resulting PELE-derived CBFE values were used as the final structural criterion for candidate ranking prior to experimental characterization. To assess enrichment by ML-guided prioritization, the CBFE distribution of ML-prioritized candidates was compared with that of an equivalently sized randomly selected control set following PELE evaluation (Figure 6).

**Figure 6.**
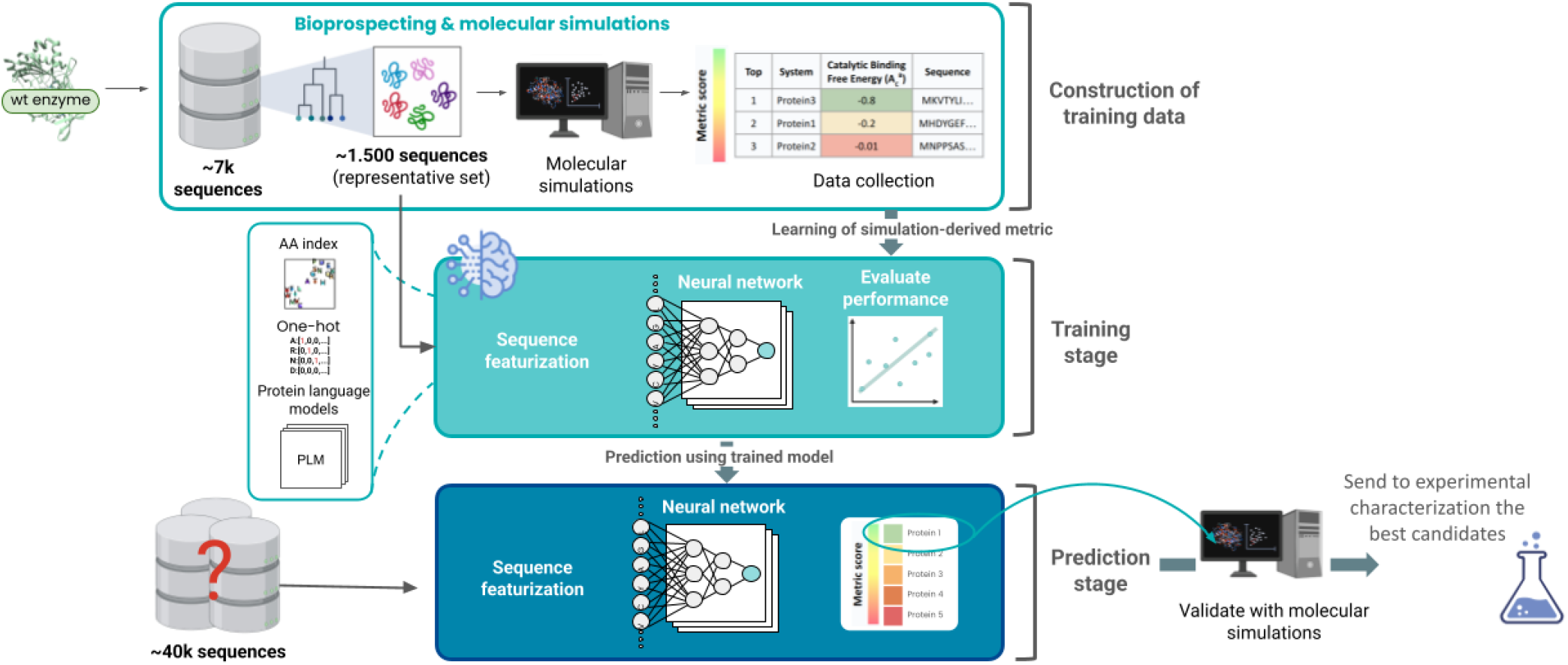
Overview of the machine-learning-guided protein sequence prioritization workflow Starting from a wild-type enzyme, approximately 7,000 sequences were found through bioprospecting, yielding -1.500 candidates for molecular simulations after filtering. Simulation-derived activity metrics were used to construct a training dataset, which was combined with sequence representations derived from amino-acid indices and pretrained protein language models. A neural network was trained to learn the relationship between sequence features and the simulation-derived metric, with model performance evaluated on held-out data. The trained model was subsequently applied to a larger sequence space to predict and rank candidates. Top-ranked sequences were further evaluated using molecular simulations before selection for experimental characterization.

## Supplementary Information

**Figure S1.**
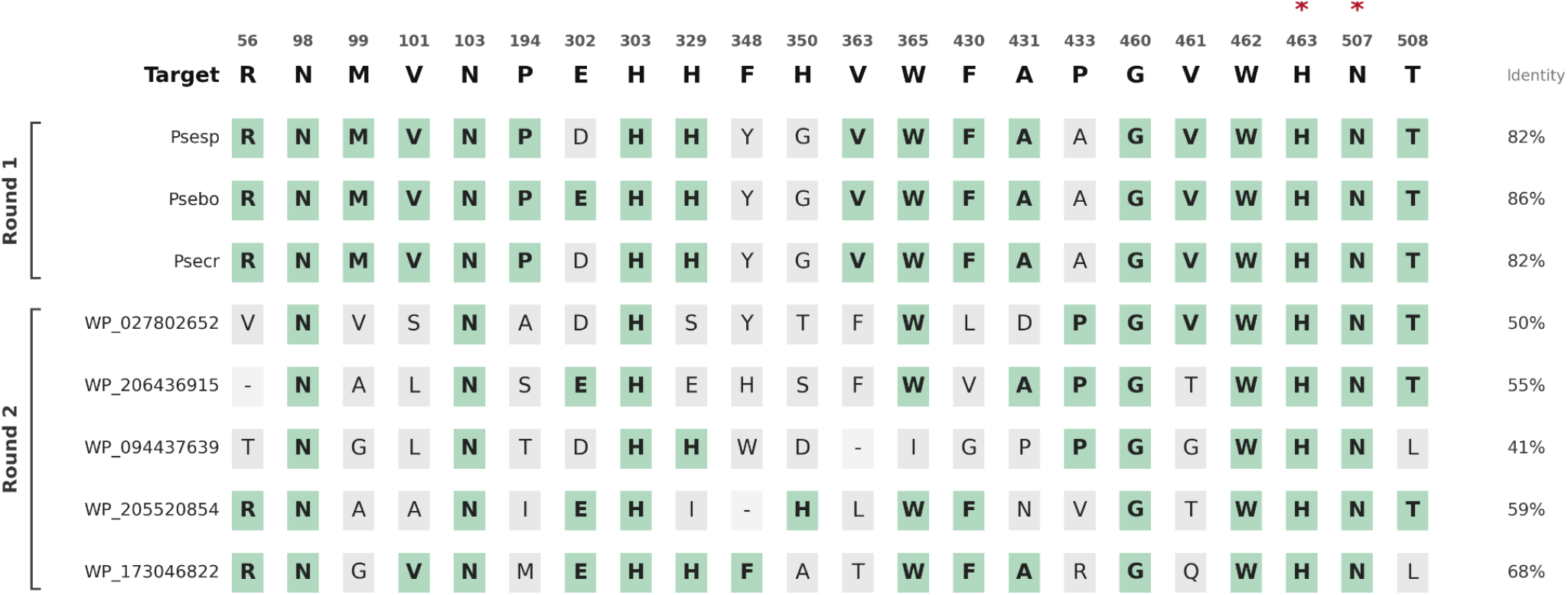
Multialignment of only the active site residues comparing round 1 and 2 with the target sequence (HMFO from Methylovorus sp. strain MP688). Residues matching the target sequence are colored in green, and the identity of this subset of residues is shown at the right of each sequence. Catalytic residues are marked with a red asterisk.

